# Novel TREM2 splicing isoform that lacks the V-set immunoglobulin domain is abundant in the human brain

**DOI:** 10.1101/2020.11.30.404897

**Authors:** Kostantin Kiianitsa, Irina Kurtz, Neal Beeman, Mark Matsushita, Wei-Ming Chien, Wendy H. Raskind, Olena Korvatska

**Affiliations:** Department of Immunology, University of Washington, Seattle, USA; Department of Medicine, Division of Medical Genetics, University of Washington, Seattle, USA; Department of Psychiatry and Behavioral Sciences, University of Washington, Seattle, USA; Mental Illness Research, Education and Clinical Center (MIRECC), VA Puget Sound Medical Center, Seattle, USA; Geriatric Research, Education and Clinical Center (GRECC), VA Puget Sound Medical Center, Seattle, USA

## Abstract

TREM2 is an immunoglobulin-like receptor expressed by certain myeloid cells, such as macrophages, dendritic cells, osteoclasts and microglia. In the brain, TREM2 plays an important role in the immune function of microglia, and its dysfunction is linked to various neurodegenerative conditions in humans. Ablation of TREM2 or its adaptor protein TYROBP causes Polycystic Lipo-Membranous Osteodysplasia with Sclerosing Leukoencephalopathy (also known as Nasu-Hakola disorder) with early onset of dementia, while some missense variants in TREM2 are associated with an increased risk of late-onset Alzheimer’s disease. Human TREM2 gene is a subject to alternative splicing, and its major, full-length canonical transcript encompasses 5 exons. Herein, we report a novel alternatively spliced TREM2 isoform without exon 2 (Δe2), which constitutes a sizable fraction of TREM2 transcripts and has highly variable inter-individual expression in the human brain (average frequency 10%; range 3.7-35%). The protein encoded by Δe2 lacks a V-set immunoglobulin domain from its extracellular part but retains its transmembrane and cytoplasmic domains. We demonstrated Δe2 protein expression in TREM2-positive THP-1 cells, in which the expression of full-length transcript was precluded by CRISPR/Cas9 disruption of the exon 2 coding frame. In “add-back” experiments, overexpression of full-length, but not Δe2 TREM2, restored phagocytic capacity and promoted interferon type I response in the knockout cells. Our findings suggest that expression of a Δe2 splice isoform may modify the dosage of full-length transcript potentially weakening some TREM2 receptor functions in the human brain.

## Introduction

**T**riggering **r**eceptor **e**xpressed on **m**yeloid cells **2** (TREM2) is an immunoglobulin-like molecule expressed by some myeloid cells, such as macrophages, dendritic cells, osteoclasts and microglia. It is an important regulator of the innate immune response and its deficiency is linked to several neurodegenerative disorders in humans (1). TREM2 forms a signaling receptor complex with TYROBP adaptor protein. Loss-of-function (LOF) bi-allelic mutations in TREM2 or TYROBP cause Polycystic Lipo-Membranous Osteodysplasia with Sclerosing Leukoencephalopathy (PLOSL; OMIM # 618193 and # 221770), a recessive disorder that presents with early onset dementia and bone cysts (2), or familial Frontotemporal Dementia without bone cysts (FTD) (3),(4),(5). PLOSL patients appear cognitively and neurologically normal till the 4th decade, highlighting a neuroprotective role of TREM2 in adult/aging microglia. The TREM2 missense variants R47H and R62H are associated with an increased risk of Alzheimer’s disease (AD) (OR 2.8-4.6 (6),(7) and 1.4-2.4 (8),(9), respectively), and R47H may also be linked to Parkinson’s disorder (10) and Amyotrophic Lateral Sclerosis (11).

TREM2 is involved in multiple aspects of myeloid cell function (12). In microglia, it regulates chemotaxis, enhances phagocytosis (13) and influences survival, proliferation and differentiation (14). Recently, we demonstrated a novel role of TREM2 in the regulation of antiviral interferon type I response (IFN I) in myeloid cells and exaggerated response in the brain of R47H carriers with AD (15). TREM2 binds lipids and lipoproteins (16), including the known AD risk factors ApoE and ApoJ/CLU (17). Another TREM2 ligand is amyloid beta peptide (Aβ) (18), a major constituent of amyloid deposits in AD brains. TREM2 binding to Aβ activates microglia cytokine production and degradation of internalized peptide. A soluble form of TREM2, a product of cleavage or alternative splicing, may have a separate function as an extracellular signaling molecule that promotes cell survival (19).

The human TREM2 gene is represented by several alternatively spliced transcripts (Fig. 1A). A major canonical form, ENST00000373113, comprises 5 exons and encodes the full-length transmembrane receptor protein. Two minor isoforms are products of alternative splicing of exon 4 that encodes the transmembrane domain. In ENST00000338469, exon 4 is skipped, while in ENST00000373122 exon 4 has an alternative start that changes its coding sequence. As a result, both isoforms lack the transmembrane domain and are thought to be secreted proteins. All three TREM2 isoforms are expressed in the human cortex (8). Herein, we report a novel alternatively spliced TREM2 transcript lacking exon 2 which constitutes a sizable fraction of TREM2 transcripts in the brain and has functional activities different from the canonical TREM2 receptor.

**Figure 1.**
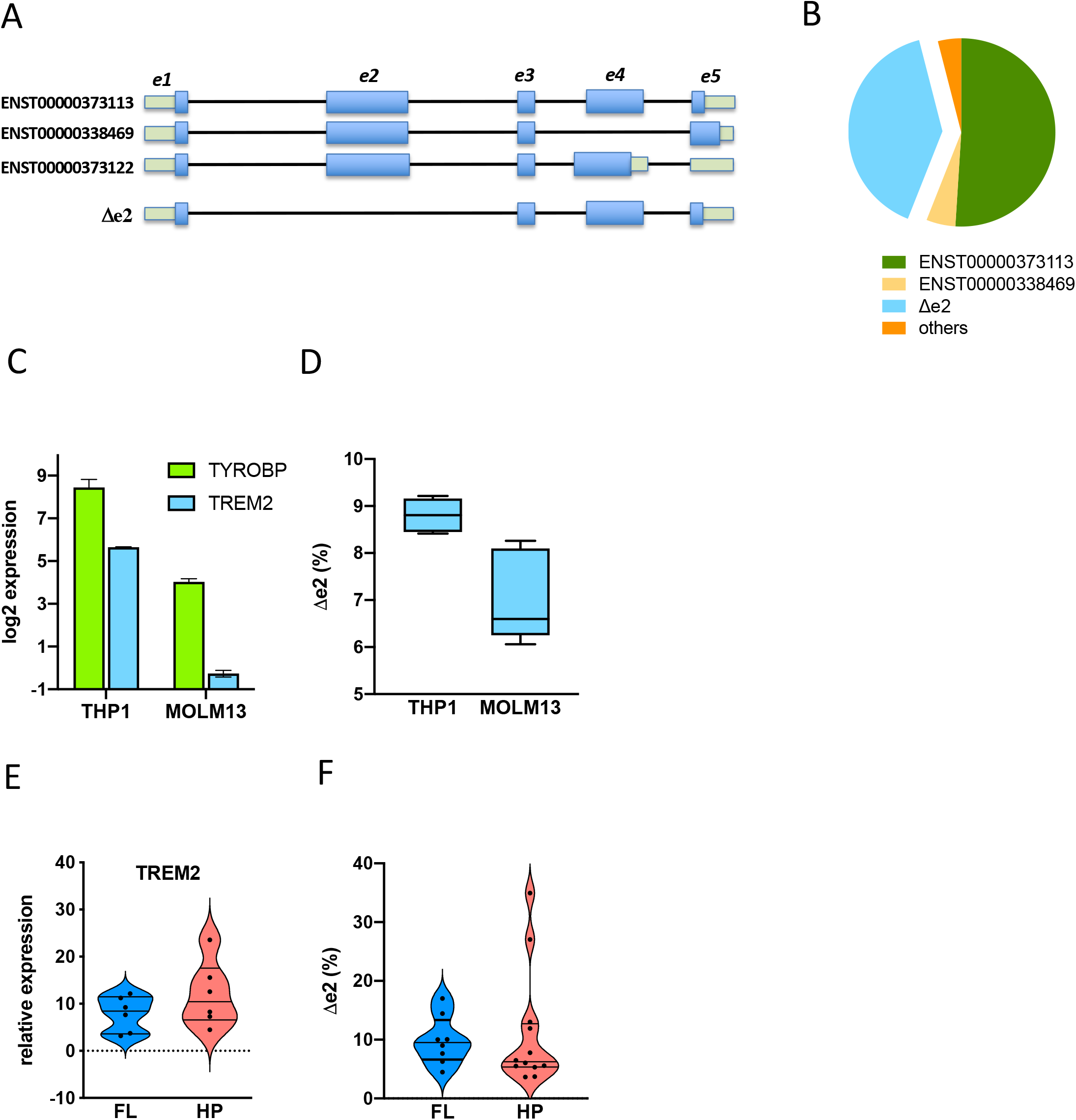
Δe2 TREM2 splice isoform is enriched in human brain. (**A**) Δe2 and three annotated (canonical) alternatively spliced TREM2 transcripts that retained exon 2. Blue boxes mark coding sequence in exons. (**B**) Frequencies of isoforms in the cDNA library cloned from human frontal cortex (four subjects, N=75 clones sequenced). (**C**) Relative levels of TREM2 and TYROBP transcripts expressed by TREM2-positive THP-1 and MOLM-13 cell lines. (**D**) fraction of Δe2 transcript expressed by these cells. (**E**) Relative levels of TREM2 expressed in the frontal lobe (FL, N=6) and hippocampus (HP, N=6) of the human brain. Gene expression in C and E was measured using qRT-PCR with commercial TaqMan assays for TYROBP and for major TREM2 isoforms (specific for e3/e4 junction), expression levels were normalized to TBP expression. (**F**) fraction of Δe2 transcript expressed in FL (N=8) and HP (N=12). D, F - Levels of Δe2 and isoforms that retained exon 2 were measured using custom TaqMan assays specific for e1/e3 and e2/e3 junctions, respectively. Absolute copy number of each isoform was calculated using calibration curves of corresponding plasmid standards.

## Materials and Methods

### Brain tissues for RNA analyses

The study was approved by the Institutional Review Board of the University of Washington. Postmortem human brain tissues of cognitively normal individuals were provided by the UW Neuropathology Core Brain Bank. Permission was obtained from donors for brain autopsy and the use of tissues for research purposes. The average post-mortem interval was 4.2 hr (range 2.5-6 hr). Tissue samples were flash-frozen at the time of autopsy and stored at −80°C.

### Cell lines and derivatives

Human myeloid cell lines THP-1 and MOLM-13 were obtained from the ATCC. Cells were cultured in RPMI supplemented with GlutaMax and 10% fetal bovine serum. TREM2 knockouts THP-1 (KO) were engineered using CRISPR/Cas9 disruption of the coding frame in exon 2 (15). Coding sequences of human full-length TREM2 (NM_018965.3) and the Δe2 isoform were synthesized and cloned into the doxycycline-inducible lentiviral pCW57-MCS1-2A-MCS2 vector (Addgene, #71782). For “add-back” experiments, TREM2 KO cells were transduced with the corresponding full-length or Δe2 lentiviral expression constructs. After 2 weeks of puromycin selection, resistant cells were harvested; ectopic TREM2 expression was induced with doxycycline (100 ng/ml) and validated by qRT-PCR and by Western blot.

### RNA isolation and cDNA synthesis

Total RNA from cultured cells was isolated with the RNAeasy kit (Qiagen; #74106). Total RNA from cryopreserved brain autopsies was isolated using TRIZOL reagent (Invitrogen). cDNA was synthesized using SuperScript III First-Strand Synthesis kit with oligo(dT) primers (Invitrogen, #18080051).

### TREM2 cDNA cloning

Total RNA was isolated from frontal cortex of four subjects and converted to cDNA. Full coding sequence of all known TREM2 isoforms was amplified using primers positioned within non-translated 5′ and 3′ regions; forward primer: 5′- gcagttcaagggaaagacga-3′; reverse primer: 5′-tccagctaaatatgacagtcttgg-3′. PCR products were gel-purified, cloned into pCR-XL-Topo vector (Invitrogen, # K4700) and sequenced.

### qRT-PCR assays

were performed on a 7500 Real-Time PCR System (Applied Biosystems) in technical duplicates or triplicates. TBP was used as an endogenous control. The following predesigned TaqMan assays (Life Technologies) were used: TREM2 Hs01010721_m1; TYROBP Hs00182426_m1; TBP Hs00427620_m1; IFNB Hs01077958_s1. We designed a pair of custom TaqMan assays which specifically measure the Δe2 transcript (e1/e3: probe spanning exon1/3 junction) vs full-length and other transcripts that retain exon 2 (e2/e3: probe spanning exon 2/3 junction).

Oligo sequences and concentrations for e1/e3 assay were:

forward primer: 5′-TCTTGCACAAGGCACTCT −3′ (600 nM);
reverse primer: 5′- GAACCAGAGATCTCCAGCAT −3′ (600 nM);
probe: 5′- TGTCACAGACCCCCTGGATCACCG −3′ (133 nM).

Oligo sequences and concentrations for e2/e3 assay were:

forward primer: 5′- ACGCTGCGGAATCTACAA −3′ (900 nM);
reverse primer: 5′- GAACCAGAGATCTCCAGCAT −3′ (900 nM);
probe: 5′- CTGGCAGACCCCCTGGATCACCG −3′ (200 nM)

### Western blotting

50 μg of whole cell lysates were resolved on polyacrylamide gel, transferred to PVDF membrane and stained with monoclonal TREM2 Abs that recognize C-terminal epitope (Cell Signaling Technology, #91068) at 1:1,000 dilution. For gel loading controls, monoclonal β- Tubulin Abs (Cell Signaling Technology, #86298) were used at 1:1,000 dilution.

### Phagocytosis assay

*In vitro* aggregation of amyloid peptide (Aβ_1-42_, Bachem #H-1368, 100 μM in PBS) was carried out by incubation for 4 days at 37°C; insoluble pellet was precipitated and labeled by pHrodo Red according to the manufacturer’s protocol (Life Technologies, #P36600). Phagocytosis of aggregated Aβ was measured as described (15). Briefly, THP-1 monocytes were plated at 5×10^5^/ml density, TREM2 expression was induced by 100 ng/ml doxycycline overnight (16-18 hr). Cells were incubated for 3 hours at 37°C with 0.25 μg/ml Aβ-pHrodo that only yields a fluorescent signal in an acidic lysophagosome compartment, collected and washed in ice-cold PBS with 1% BSA. Uptake of Aβ-pHrodo was measured on the LSRII Flow cytometer (BD Biosciences) and data analyzed by FlowJo. 20,000 events per sample were scored, cells were gated by forward and side scatter to exclude dead ones and doublets. Phagocytosis was calculated from mean fluorescence intensity of internalized pHrodo.

### THP-1 differentiation into macrophages and stimulation of IFN I response

Differentiation to macrophages was performed as previously described (15) using 5ng/ml phorbol 12-myristate 13-acetate (PMA) for 48 hr followed by 24 hr of recovery without PMA. IFN I response was induced by high molecular weight poly(I:C) complexes with LyoVec transfection reagent (Invivogen, tlrl-piclv, 500 ng/ml), in combination with IFNβ (PeproTech, #300-02, 100 Units/ml) for 24 hr. For “add-back” experiments, TREM2 expression was induced by 100 ng/ml doxycycline 16-18 hr prior to the IFN I response stimulation.

### Statistical analysis

was performed using Prism version 8 software (GraphPad, La Jolla, CA).

## Results and Discussion

We analyzed the distribution of TREM2 isoforms in human brains using primers to 5′ and 3′ untranslated regions. RT-PCR products were cloned and sequenced. In addition to the annotated isoforms, we identified a novel splice variant lacking exon 2 (Δe2, Fig. 1A) that comprised approximately 40% of sequenced clones (Fig. 1B). Exon 2 skipping produces in-frame deletion of amino acids 14-130 corresponding to a portion of signal peptide and the entire V-set Immunoglobulin (Ig V) domain (aa 29-112).

To confirm Δe2 is a naturally occurring TREM2 isoform, we tested THP-1 and MOLM-13, two cell lines ranked as top TREM2-TYROBP expressors among 1,443 cell lines (https://genevestigator.com, Fig. 1C). Because both PCR and cloning of fragments of different length may introduce an efficiency bias and therefore cannot reliably quantify isoforms, we designed a pair of TaqMan qRT-PCR assays, in which probes spanned either exon 2/3 or exon 1/3 junctions. The e1/e3 assay measures exclusively the Δe2 while e2/e3 assay accounts for all TREM2 isoforms that retained exon 2. We used plasmids with cloned full length and Δe2 TREM2 sequences to verify assay’s specificity and to make standard curves for absolute copy number determination. The Δe2 isoform accounted for ~9% and 7% of total TREM2 transcripts in THP-1 and MOLM-13, respectively (Fig. 1D). In human brain, Δe2 had highly variable expression ranging 3.5-17% in the frontal lobe and 3.7-35% in the hippocampus (Fig. 1F). Both areas expressed comparable levels of total TREM2 transcript with higher inter-individual variability in the hippocampus (Fig. 1E).

To confirm expression of TREM2 isoforms at protein level, we used unmodified THP-1 and cells in which exon 2 coding sequence was disrupted by CRISPR/Cas9 (TREM2 KO THP-1, (15)). The editing is expected to terminate translation of all isoforms that retain exon 2 while preserving the Δe2 isoform. Cells were stimulated by IL-4 known to induce TREM2 expression in primary macrophages (20). The membrane was probed with rabbit monoclonal antibodies recognizing an epitope at C-terminus. In unmodified cells, two major bands were observed: the upper one corresponding to full-length protein and the lower one with a predicted size of C-terminal TREM2 fragment (CTF), a membrane-associated product of proteolytic receptor shedding (Fig.2A). The cleavage occurs at extracellular H157/S158 residues (21), (22). A minor band corresponding to the predicted size of Δe2 polypeptide was located above the CTF band. As expected, TREM2 KO cells did not express full-length protein. Upon IL-4 stimulation, only the Δe2 isoform was increased in TREM2 KO, whereas all TREM2 species were increased in unmodified cells.

**Figure 2.**
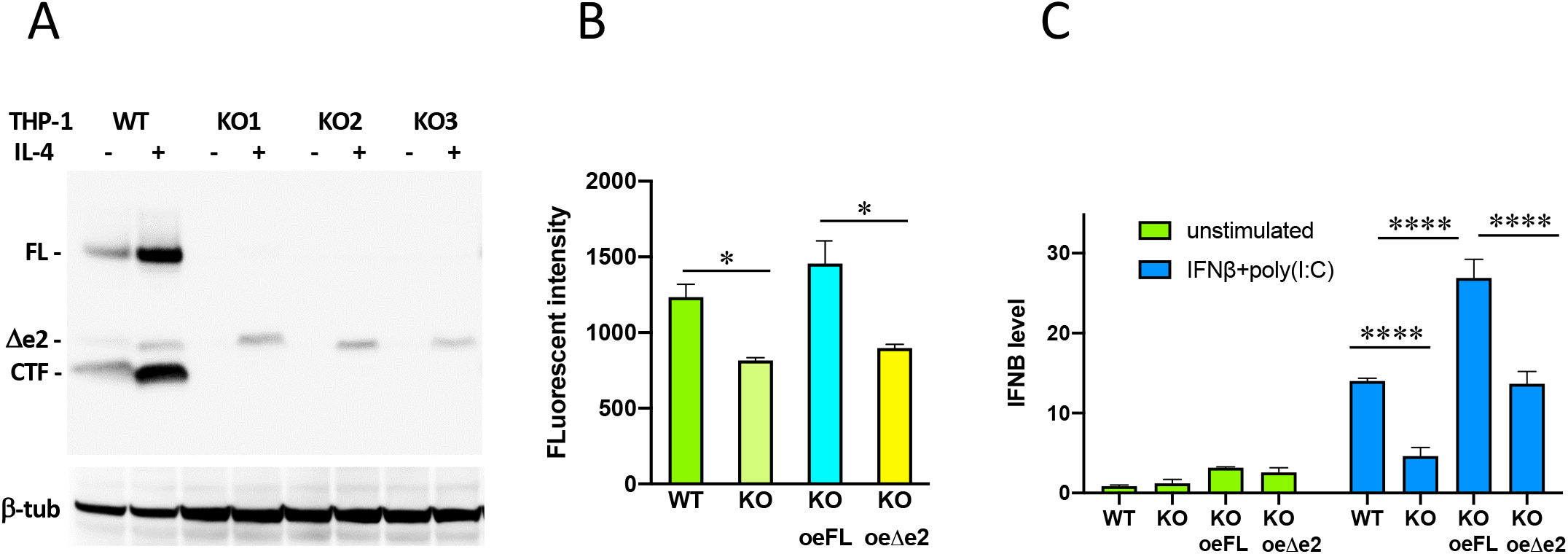
Protein expression and functional activities of Δe2 TREM2. (**A**). Western blot of THP-1 cell extracts probed with anti-TREM2 Abs, before and after 24 hr stimulation with IL-4. WT – unmodified THP-1; KO1, KO2, KO3 – independent TREM2 knockout clones. FL – full length receptor encoded by a major canonical isoform; CTF - a membrane-associated C-terminal fragment, a product of full lengths receptor shedding due to proteolysis at the amino acids 157-158 (21). Loading controls were probed with anti-tubulin Abs. (**B**). Phagocytosis of aggregated pHrodo-labeled Amyloid beta peptide (Aβ). WT – unmodified THP-1; KO - TREM2 knockout (KO1 clone); oeFL, oeΔe2 – TREM2 KO1 cells with stably integrated lentiviral doxycycline-inducible constructs expressing full length or Δe2 isoform, respectively. TREM2 overexpression was induced with doxycycline. Cells were incubated with pHrodo-labeled Aβ and fluorescence measured by flow cytometry. Shown are means ± SD of biological triplicates; * - p<0.05; one-way ANOVA (Dunnett’s multiple comparison test). (**C**). IFN I response of *in vitro* differentiated THP-1 macrophages. THP-1 monocytes were differentiated to macrophages by PMA; TREM2 expression was induced with doxycycline prior to stimulation of IFN I response with poly(I:C) + IFNβ. After 24 hr stimulation, cells were collected and total RNA isolated. IFN I response was quantified as induction of IFNB mRNA, expression levels were normalized to unstimulated unmodified THP-1. WT – unmodified THP-1; KO - TREM2 knockout (KO1 clone); oeFL, oeΔe2 – TREM2 KO1 cells with stably integrated lentiviral doxycycline-inducible constructs over-expressing full length or Δe2 isoform, respectively. Shown are means ± SD of biological triplicates; **** - p<0.0001; two-way ANOVA, (Tukey’s multiple comparison test).

We recently showed that TREM2 ablation substantially blunted IFN I response and reduced Aβ phagocytosis in THP-1 cells; these activities were restored upon overexpression of either wild type or R47H variant proteins in TREM2 KO THP-1 cells (15). To see if Δe2 isoform complements these deficits, we integrated it into TREM2 KO THP-1under control of a doxycycline-inducible promoter. Δe2 was unable to restore Aβ phagocytosis in TREM2 KO; it also had diminished ability to promote an IFN I response (Fig.2 B-C). Thus, Δe2, a naturally occurring TREM2 splice isoform enriched in the human brain, does not complement some activities of full-length TREM2.

Alternative RNA splicing is an important mechanism that generates protein diversity by reshuffling functional domains of proteins. Variants that affect splicing regulatory motifs are usually deleterious, and about one-tenth of disease-associated variants reported in the Human Gene Mutation Database are splicing mutations (23). Of known loss of function TREM2 variants responsible for PLOSL and FTD about 20% disrupt the coding frame via altered splicing. These include variants that cause intron 2 retention (24) and exon 3 skipping (25) in FTD or PLOSL patients. A variant in intron 1, c.40+3delAGG, responsible for FTD without bone cysts, weakens the donor splice site resulting in 2-fold reduction of TREM2 level (26).

Exon 2 encodes most of the extracellular TREM2 moiety harboring the AD-associated R47 and R62, as well as the majority of residues mutated in PLOSL and FTD (1). Its absence is likely to affect TREM2 interactions with known ligands and is a plausible cause of Δe2’s inability to complement the Aβ phagocytosis deficiency of TREM2 KO cells. Because exon skipping/inclusion are mutually exclusive events, production of the Δe2 transcript is expected to decrease the levels of canonical transcripts that retain exon 2. Of note, the Δe2 retains transmembrane and cytoplasmic domains, essentially mimicking the CTF, a membrane-associated product of receptor shedding. Future studies will be required to determine whether Δe2 has an additional regulatory role, for instance by competing for binding with the TYROBP adaptor and sequestering full length TREM2 from the receptor complex.

TREM2 resides within the TREM gene cluster on chromosome 6p21.1, which also includes structurally similar paralogs TREM1, TREML1, TREML2 and TREML4. Coding and non-coding variants in these genes were found to modify AD risks, independently of TREM2 R47H (27), (28). Interestingly, the TREML1/TLT-1 paralog also has an alternatively spliced transcript (ENST00000437044.2) (29) with skipped exon 2 corresponding to an entire Ig V domain. This suggests that Ig V inclusion/exclusion via alternative splicing is one of nature’s tool for functional diversification of some of the Ig-like immune molecules.

The inhibitory CD33 receptor is another microglia-specific AD risk gene (30), (31). Similar to TREM2, CD33 belongs to the superfamily of Ig-like immune receptors, and its Ig V domain is encoded by exon 2 (Fig. 3). Intriguingly, the AD-associated CD33 risk allele, rs3865444(C), instructs inclusion of exon 2 and production of full-length protein, while a protective allele, rs3865444(A), instructs exon 2 skipping and production of Δe2 CD33 (32), (33). The full-length CD33 exerts an inhibitory effect on some microglia activities, such as cytokine production and/or phagocytosis in response to AD-relevant stimuli, whereas Δe2 CD33 is unable to suppress microglia activation and amyloid plaque phagocytosis *in vitro* and *in vivo* (34). While the regulatory variants that change AD risk by instructing inclusion/exclusion of Ig V-encoding exon 2 are known for CD33, the impact of cis-regulatory variation on TREM2 splicing and its contribution to neurodegenerative pathologies remain to be elucidated.

**Figure 3.**
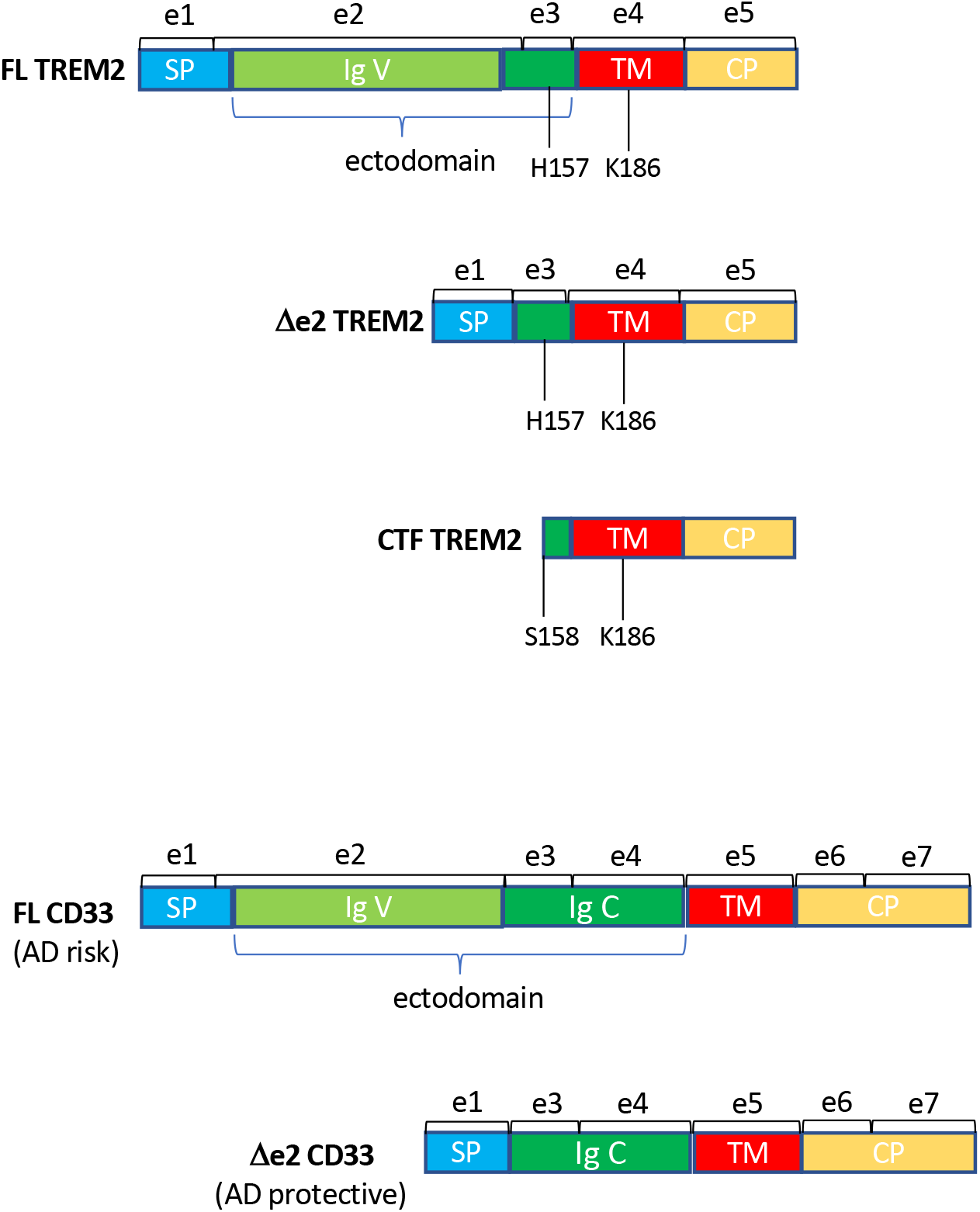
Domain organization of isoforms of microglial receptors TREM2 and CD33. SP - signal peptide; TM - transmembrane domain; CP - cytoplasmic domain; Ig V, Ig C - Immunoglobulin-like domains; FL - full length; CTF - C-terminal fragment, Δe2 - lacking exon 2. H157/S158 - proteolytic shedding site; K186 - conserved residue responsible for interaction with TYROBP adaptor. Corresponding exons are depicted by square brackets. The diagram is not drawn to scale.

## AUTHORSHIP

K. K. designed and performed the experiments, analyzed the results, and wrote the manuscript, I. K., M.M., N. B., N.L. and W-M. C. performed the experiments and analyzed the results; W. H. R. and O.K. supervised the studies, analyzed and interpreted the results, wrote the manuscript and acquired funding.

## ACKNOWLEDGMENTS

This work was supported by National Institute of Health grants [P30AG013280 to O.K., 2R01 NS069719 to W.H.R. and Merit Review Award Number 101 CX001702 from the United States (U.S.) Department of Veterans Affairs Clinical Sciences R&D (CSRD) Service to W.H.R.]

## References

1 Ulland, T. K. and Colonna, M. 2018. TREM2 - a key player in microglial biology and Alzheimer disease. Nature reviews. Neurology 14:667.

2 Paloneva, J., Manninen, T., Christman, G., Hovanes, K., Mandelin, J., Adolfsson, R., Bianchin, M., Bird, T., Miranda, R., Salmaggi, A., Tranebjaerg, L., Konttinen, Y., and Peltonen, L. 2002. Mutations in two genes encoding different subunits of a receptor signaling complex result in an identical disease phenotype. American journal of human genetics 71:656.

3 Guerreiro, R., Bilgic, B., Guven, G., Bras, J., Rohrer, J., Lohmann, E., Hanagasi, H., Gurvit, H., and Emre, M. 2013. Novel compound heterozygous mutation in TREM2 found in a Turkish frontotemporal dementia-like family. Neurobiology of aging 34:2890 e1.

4 Guerreiro, R. J., Lohmann, E., Bras, J. M., Gibbs, J. R., Rohrer, J. D., Gurunlian, N., Dursun, B., Bilgic, B., Hanagasi, H., Gurvit, H., Emre, M., Singleton, A., and Hardy, J. 2013. Using exome sequencing to reveal mutations in TREM2 presenting as a frontotemporal dementia-like syndrome without bone involvement. JAMA neurology 70:78.

5 Le Ber, I., De Septenville, A., Guerreiro, R., Bras, J., Camuzat, A., Caroppo, P., Lattante, S., Couarch, P., Kabashi, E., Bouya-Ahmed, K., Dubois, B., and Brice, A. 2014. Homozygous TREM2 mutation in a family with atypical frontotemporal dementia. Neurobiology of aging 35:2419 e23.

6 Guerreiro, R., Wojtas, A., Bras, J., Carrasquillo, M., Rogaeva, E., Majounie, E., Cruchaga, C., Sassi, C., Kauwe, J. S., Younkin, S., Hazrati, L., Collinge, J., Pocock, J., Lashley, T., Williams, J., Lambert, J. C., Amouyel, P., Goate, A., Rademakers, R., Morgan, K., Powell, J., St George-Hyslop, P., Singleton, A., and Hardy, J. 2012. TREM2 Variants in Alzheimer’s Disease. The New England journal of medicine.

7 Jonsson, T., Stefansson, H., Ph, D. S., Jonsdottir, I., Jonsson, P. V., Snaedal, J., Bjornsson, S., Huttenlocher, J., Levey, A. I., Lah, J. J., Rujescu, D., Hampel, H., Giegling, I., Andreassen, O. A., Engedal, K., Ulstein, I., Djurovic, S., Ibrahim-Verbaas, C., Hofman, A., Ikram, M. A., van Duijn, C. M., Thorsteinsdottir, U., Kong, A., and Stefansson, K. 2012. Variant of TREM2 Associated with the Risk of Alzheimer’s Disease. The New England journal of medicine.

8 Jin, S. C., Benitez, B. A., Karch, C. M., Cooper, B., Skorupa, T., Carrell, D., Norton, J. B., Hsu, S., Harari, O., Cai, Y., Bertelsen, S., Goate, A. M., and Cruchaga, C. 2014. Coding variants in TREM2 increase risk for Alzheimer’s disease. Human molecular genetics 23:5838.

9 Song, W., Hooli, B., Mullin, K., Jin, S. C., Cella, M., Ulland, T. K., Wang, Y., Tanzi, R. E., and Colonna, M. 2017. Alzheimer’s disease-associated TREM2 variants exhibit either decreased or increased ligand-dependent activation. Alzheimer’s & dementia : the journal of the Alzheimer’s Association 13:381.

10 Rayaprolu, S., Mullen, B., Baker, M., Lynch, T., Finger, E., Seeley, W. W., Hatanpaa, K. J., Lomen-Hoerth, C., Kertesz, A., Bigio, E. H., Lippa, C., Josephs, K. A., Knopman, D. S., White, C. L., 3rd, Caselli, R., Mackenzie, I. R., Miller, B. L., Boczarska-Jedynak, M., Opala, G., Krygowska-Wajs, A., Barcikowska, M., Younkin, S. G., Petersen, R. C., Ertekin-Taner, N., Uitti, R. J., Meschia, J. F., Boylan, K. B., Boeve, B. F., Graff-Radford, N. R., Wszolek, Z. K., Dickson, D. W., Rademakers, R., and Ross, O. A. 2013. TREM2 in neurodegeneration: evidence for association of the p.R47H variant with frontotemporal dementia and Parkinson’s disease. Molecular neurodegeneration 8:19.

11 Cady, J., Koval, E. D., Benitez, B. A., Zaidman, C., Jockel-Balsarotti, J., Allred, P., Baloh, R. H., Ravits, J., Simpson, E., Appel, S. H., Pestronk, A., Goate, A. M., Miller, T. M., Cruchaga, C., and Harms, M. B. 2014. TREM2 variant p.R47H as a risk factor for sporadic amyotrophic lateral sclerosis. JAMA neurology 71:449.

12 Deczkowska, A., Weiner, A., and Amit, I. 2020. The Physiology, Pathology, and Potential Therapeutic Applications of the TREM2 Signaling Pathway. Cell 181:1207.

13 Kleinberger, G., Yamanishi, Y., Suarez-Calvet, M., Czirr, E., Lohmann, E., Cuyvers, E., Struyfs, H., Pettkus, N., Wenninger-Weinzierl, A., Mazaheri, F., Tahirovic, S., Lleo, A., Alcolea, D., Fortea, J., Willem, M., Lammich, S., Molinuevo, J. L., Sanchez-Valle, R., Antonell, A., Ramirez, A., Heneka, M. T., Sleegers, K., van der Zee, J., Martin, J. J., Engelborghs, S., Demirtas-Tatlidede, A., Zetterberg, H., Van Broeckhoven, C., Gurvit, H., Wyss-Coray, T., Hardy, J., Colonna, M., and Haass, C. 2014. TREM2 mutations implicated in neurodegeneration impair cell surface transport and phagocytosis. Science translational medicine 6:243ra86.

14 Zheng, H., Jia, L., Liu, C. C., Rong, Z., Zhong, L., Yang, L., Chen, X. F., Fryer, J. D., Wang, X., Zhang, Y. W., Xu, H., and Bu, G. 2017. TREM2 Promotes Microglial Survival by Activating Wnt/beta-Catenin Pathway. The Journal of neuroscience : the official journal of the Society for Neuroscience 37:1772.

15 Korvatska, O., Kiianitsa, K., Ratushny, A., Matsushita, M., Beeman, N., Chien, W. M., Satoh, J. I., Dorschner, M. O., Keene, C. D., Bammler, T., Bird, T. D., and Raskind, W. H. 2020. TREM2 R47H exacerbates immune response in Alzheimer’s disease brain. Front Immunol 11.

16 Wang, Y., Cella, M., Mallinson, K., Ulrich, J. D., Young, K. L., Robinette, M. L., Gilfillan, S., Krishnan, G. M., Sudhakar, S., Zinselmeyer, B. H., Holtzman, D. M., Cirrito, J. R., and Colonna, M. 2015. TREM2 lipid sensing sustains the microglial response in an Alzheimer’s disease model. Cell 160:1061.

17 Yeh, F. L., Wang, Y., Tom, I., Gonzalez, L. C., and Sheng, M. 2016. TREM2 Binds to Apolipoproteins, Including APOE and CLU/APOJ, and Thereby Facilitates Uptake of Amyloid-Beta by Microglia. Neuron 91:328.

18 Zhao, Y., Wu, X., Li, X., Jiang, L. L., Gui, X., Liu, Y., Sun, Y., Zhu, B., Pina-Crespo, J. C., Zhang, M., Zhang, N., Chen, X., Bu, G., An, Z., Huang, T. Y., and Xu, H. 2018. TREM2 Is a Receptor for beta-Amyloid that Mediates Microglial Function. Neuron 97:1023.

19 Zhong, L., Chen, X. F., Wang, T., Wang, Z., Liao, C., Wang, Z., Huang, R., Wang, D., Li, X., Wu, L., Jia, L., Zheng, H., Painter, M., Atagi, Y., Liu, C. C., Zhang, Y. W., Fryer, J. D., Xu, H., and Bu, G. 2017. Soluble TREM2 induces inflammatory responses and enhances microglial survival. The Journal of experimental medicine 214:597.

20 Turnbull, I. R., Gilfillan, S., Cella, M., Aoshi, T., Miller, M., Piccio, L., Hernandez, M., and Colonna, M. 2006. Cutting edge: TREM-2 attenuates macrophage activation. J Immunol 177:3520.

21 Thornton, P., Sevalle, J., Deery, M. J., Fraser, G., Zhou, Y., Stahl, S., Franssen, E. H., Dodd, R. B., Qamar, S., Gomez Perez-Nievas, B., Nicol, L. S., Eketjall, S., Revell, J., Jones, C., Billinton, A., St George-Hyslop, P. H., Chessell, I., and Crowther, D. C. 2017. TREM2 shedding by cleavage at the H157-S158 bond is accelerated for the Alzheimer’s disease-associated H157Y variant. EMBO Mol Med 9:1366.

22 Schlepckow, K., Kleinberger, G., Fukumori, A., Feederle, R., Lichtenthaler, S. F., Steiner, H., and Haass, C. 2017. An Alzheimer-associated TREM2 variant occurs at the ADAM cleavage site and affects shedding and phagocytic function. EMBO Mol Med 9:1356.

23 Stenson, P. D., Mort, M., Ball, E. V., Evans, K., Hayden, M., Heywood, S., Hussain, M., Phillips, A. D., and Cooper, D. N. 2017. The Human Gene Mutation Database: towards a comprehensive repository of inherited mutation data for medical research, genetic diagnosis and next-generation sequencing studies. Human genetics 136:665.

24 Li, X., Sun, Y., Gong, L., Zheng, L., Chen, K., Zhou, Y., Gu, Y., Xu, Y., Guo, Q., Hong, Z., Ding, D., Fu, J., and Zhao, Q. 2019. A novel homozygous mutation in TREM2 found in a Chinese early-onset dementia family with mild bone involvement. Neurobiology of aging.

25 Numasawa, Y., Yamaura, C., Ishihara, S., Shintani, S., Yamazaki, M., Tabunoki, H., and Satoh, J. I. 2011. Nasu-Hakola disease with a splicing mutation of TREM2 in a Japanese family. European journal of neurology : the official journal of the European Federation of Neurological Societies 18:1179.

26 Chouery, E., Delague, V., Bergougnoux, A., Koussa, S., Serre, J. L., and Megarbane, A. 2008. Mutations in TREM2 lead to pure early-onset dementia without bone cysts. Human mutation 29:E194.

27 Benitez, B. A., Jin, S. C., Guerreiro, R., Graham, R., Lord, J., Harold, D., Sims, R., Lambert, J. C., Gibbs, J. R., Bras, J., Sassi, C., Harari, O., Bertelsen, S., Lupton, M. K., Powell, J., Bellenguez, C., Brown, K., Medway, C., Haddick, P. C., van der Brug, M. P., Bhangale, T., Ortmann, W., Behrens, T., Mayeux, R., Pericak-Vance, M. A., Farrer, L. A., Schellenberg, G. D., Haines, J. L., Turton, J., Braae, A., Barber, I., Fagan, A. M., Holtzman, D. M., Morris, J. C., Group, C. S., consortium, E., Alzheimer’s Disease Genetic, C., Alzheimer’s Disease Neuroimaging, I., Consortium, G., Williams, J., Kauwe, J. S., Amouyel, P., Morgan, K., Singleton, A., Hardy, J., Goate, A. M., and Cruchaga, C. 2014. Missense variant in TREML2 protects against Alzheimer’s disease. Neurobiology of aging 35:1510 e19.

28 Carrasquillo, M. M., Allen, M., Burgess, J. D., Wang, X., Strickland, S. L., Aryal, S., Siuda, J., Kachadoorian, M. L., Medway, C., Younkin, C. S., Nair, A., Wang, C., Chanana, P., Serie, D., Nguyen, T., Lincoln, S., Malphrus, K. G., Morgan, K., Golde, T. E., Price, N. D., White, C. C., De Jager, P. L., Bennett, D. A., Asmann, Y. W., Crook, J. E., Petersen, R. C., Graff-Radford, N. R., Dickson, D. W., Younkin, S. G., and Ertekin-Taner, N. 2017. A candidate regulatory variant at the TREM gene cluster associates with decreased Alzheimer’s disease risk and increased TREML1 and TREM2 brain gene expression. Alzheimer’s & dementia : the journal of the Alzheimer’s Association 13:663.

29 Gerhard, D. S., Wagner, L., Feingold, E. A., Shenmen, C. M., Grouse, L. H., et al. 2004. The status, quality, and expansion of the NIH full-length cDNA project: the Mammalian Gene Collection (MGC). Genome research 14:2121.

30 Carrasquillo, M. M., Belbin, O., Hunter, T. A., Ma, L., Bisceglio, G. D., Zou, F., Crook, J. E., Pankratz, V. S., Sando, S. B., Aasly, J. O., Barcikowska, M., Wszolek, Z. K., Dickson, D. W., Graff-Radford, N. R., Petersen, R. C., Passmore, P., Morgan, K., Alzheimer’s Research, U. K. c., and Younkin, S. G. 2011. Replication of EPHA1 and CD33 associations with late-onset Alzheimer’s disease: a multi-centre case-control study. Molecular neurodegeneration 6:54.

31 Bradshaw, E. M., Chibnik, L. B., Keenan, B. T., Ottoboni, L., Raj, T., Tang, A., Rosenkrantz, L. L., Imboywa, S., Lee, M., Von Korff, A., Alzheimer Disease Neuroimaging, I., Morris, M. C., Evans, D. A., Johnson, K., Sperling, R. A., Schneider, J. A., Bennett, D. A., and De Jager, P. L. 2013. CD33 Alzheimer’s disease locus: altered monocyte function and amyloid biology. Nat Neurosci 16:848.

32 Malik, M., Simpson, J. F., Parikh, I., Wilfred, B. R., Fardo, D. W., Nelson, P. T., and Estus, S. 2013. CD33 Alzheimer’s risk-altering polymorphism, CD33 expression, and exon 2 splicing. The Journal of neuroscience : the official journal of the Society for Neuroscience 33:13320.

33 Raj, T., Ryan, K. J., Replogle, J. M., Chibnik, L. B., Rosenkrantz, L., Tang, A., Rothamel, K., Stranger, B. E., Bennett, D. A., Evans, D. A., De Jager, P. L., and Bradshaw, E. M. 2014. CD33: increased inclusion of exon 2 implicates the Ig V-set domain in Alzheimer’s disease susceptibility. Human molecular genetics 23:2729.

34 Griciuc, A., Serrano-Pozo, A., Parrado, A. R., Lesinski, A. N., Asselin, C. N., Mullin, K., Hooli, B., Choi, S. H., Hyman, B. T., and Tanzi, R. E. 2013. Alzheimer’s disease risk gene CD33 inhibits microglial uptake of amyloid beta. Neuron 78:631.

